# Gene replacement therapy for Piga GPI-anchor deficiency in the developing nervous system

**DOI:** 10.64898/2025.12.11.693709

**Authors:** Jennifer L. Watts, Shibi Likhite, Yoshiko Murakami, Taroh Kinoshita, Kathrin Meyer, Rolf W. Stottmann

**Author notes:** Correspondence: Rolf W. Stottmann, Institute for Genomic Medicine, Nationwide Children’s Hospital, 575 Children’s Crossroad, Columbus, OH 43215.

## Abstract

Glycosylphosphatidylinositol (GPI) anchors are a class of post-translational modifications observed on over 150 proteins. Pathogenic variants in the GPI biosynthesis enzyme, *PIGA*, in humans are associated with several brain anomalies such as hypomyelination, cerebellar hypoplasia, ataxic gait, and can lead to premature mortality. We previously genetically deleted *Piga* from the embryonic mouse brain which led to early postnatal death and significant structural brain malformations similar to those observed in humans with *PIGA* variants. The current treatment options for *PIGA* patients only manage symptoms and provides palliative care, demonstrating a need for new therapeutic options. We employed an AAV9-mediated *PIGA* gene-replacement (AAV9-*hPIGA*) strategy to assess the efficacy of gene therapy in the brain. We show that a single intracerebroventricular treatment on the first day of life successfully rescued survival rates, structural brain anomalies, and neurological impairments. Additionally, we used mass spectrometry to identify and quantify GPI-anchored proteins in untreated and treated mutant mice. We found that AAV9- *hPIGA* treatment restored GPI-anchored protein levels in mutant animals. These investigations enhance our understanding of GPI-anchored protein production during brain development and contribute to the development of a more effective intervention for *PIGA*-related symptoms.

**Graphical Abstract:** 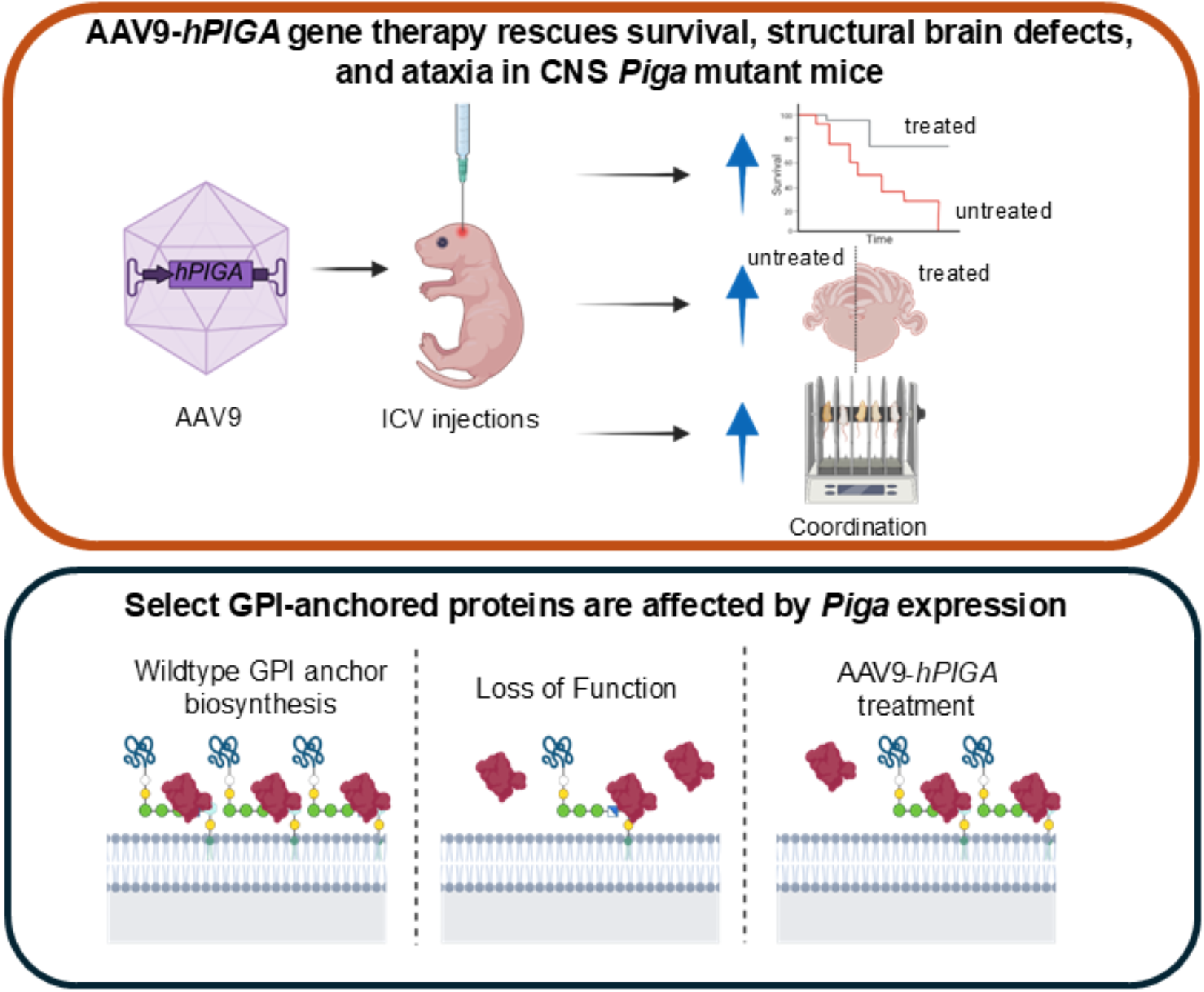

**Key Points:**

- A single AAV9-*hPIGA* injection rescues *Piga*-related phenotypes including survival and structural brain defects
- AAV9-*hPIGA* treatment rescued long-term ataxia phenotypes in *Piga* mutants
- Neuronal GPI-anchored protein levels in *Piga* mutants are increased with AAV9- *hPIGA* gene replacement

## Introduction

Posttranslational modifications of proteins are often essential for proper function and molecular interactions. Glycosylphosphatidylinositol (GPI) anchors are post-translational modifications that attach to more than 150 proteins to function as adhesion molecules, membrane enzymes and receptors (1-3). Some of these 150 proteins include CONTACTIN1, EFNA3, and GPC1, which are important for neurogenesis (4). GPI-anchor biosynthesis utilizes a cascade of twenty-eight enzymes starting with Phosphatidylinositol Glycan Biosynthesis Class (PIG) proteins in the endoplasmic reticulum (2, 5-9). Phosphatidylinositol Glycan Biosynthesis Class A (PIGA) is the first catalytic enzyme of the glycosylphosphatidylinositol-N-acetylglucosaminyltransferase complex and begins GPI-anchor biosynthesis. The complex attaches a Glc-NAC phosphatidylinositol to target proteins on the cytoplasmic side of the endoplasmic reticulum.

Inherited GPI Deficiencies are caused by pathogenic variants in the genes encoding GPI-anchor biosynthesis enzymes that are associated with several congenital disorders. These conditions show a wide variety of neural and craniofacial phenotypes including developmental delay, epilepsy, cleft lip/palate, dysmorphic features, and a considerable number infants die prematurely (10). Other phenotypes include delayed white matter development and/or atrophy, thin corpus callosum, cerebellar hypoplasia and/or atrophy, and microcephaly (11-32). *PIGA*, located on the X chromosome, causes an X-linked recessive disorder with pathogenic variants. Due to skewed X-inactivation, female carriers exhibit no phenotypes, while males carrying pathogenic variants are affected. Current treatment options are mostly limited to managing symptoms (10, 21, 29, 33) and therefore, the development of new treatment strategies is essential to better treat patients with these disorders.

Advances in adeno-associated virus (AAV) technology have established a platform for delivering long-term gene therapy to cells. AAVs can carry 4.7 kilobases of genetic material to stimulate gene expression or perform CRISPR gene editing, among other uses (34). The rapid development of this technology has also met with clinical success in treating various disorders, including cancers and neurological and neuromuscular conditions (35-38). This success is mainly due to engineering different AAV vectors to improve tissue and cell type targeting using a combination of various AAV serotypes to take advantage of different tissue tropisms and/or using specific promoters to stimulate cell type specific expression of AAV delivered constructs (39). Mouse and cellular models of inherited GPI deficiency have been used for AAV-mediated preclinical gene therapy trials for *Piga* and *Pigo* mutations, and have led to improvements in central nervous system phenotypes (40, 41).

We previously published a genetic ablation of *Piga* in the central nervous system (*Nestin-Cre; Piga*) which exhibited many phenotypes reminiscent of patients with *PIGA* pathogenic variants. These included abnormal cerebellum, ataxia, hypomyelination, and premature death (42). The pattern of inheritance in affected patients with inherited GPI deficiency and molecular data in the literature suggest that the disease results from partial loss-of-function variants, making gene replacement therapy an attractive intervention strategy. Adeno-associated viral vector type 9 (AAV9)-mediated gene therapy has shown great promise in treating CNS disorders (43-46). Here, we leverage the *Nestin-Cre; Piga* mouse model with robust CNS phenotypes to also demonstrate the power of AAV9-mediated rescue of GPI-anchor deficiency phenotypes. Additionally, we identify the specific neural GPI-anchored proteins affected by *Piga* expression using a proteomic approach to further understand the potential mechanisms underlying *Piga* phenotypes and *Piga*-mediated neurodevelopment.

## Results

### AAV9-hPIGA gene replacement therapy rescues Piga-associated lethality and brain defects

AAV-based gene replacement therapies are robust treatments in preclinical models and in the clinic for neurological and neuromuscular disorders (35-37, 40, 44-60). The *Nestin-Cre; Piga* mouse models (*Nestin*^*Cre/wt*^; *Piga*^*flox/X*^ females and *Nestin*^*Cre/wt*^; *Piga*^*flox/Y*^ males) recapitulate many of the neurological symptoms seen in human *PIGA* patients (42). Therefore, we used this mouse model to assess the efficacy of a gene-replacement therapy strategy. We generated two viral vectors designed to express human *PIGA* or *GFP* under control of the strong CMV enhancer/chicken-beta actin promoter (CBA; scAAV9.CBA.*hPIGA*, referred to as AAV9-*hPIGA* and scAAV9.CBA.GFP referred to as AAV9-*GFP*). We administered a single dose of AAV9-*hPIGA* (either 5e10 viral genomes (vg) or 7e10 vg) in a 5 mL injection volume by intracerebroventricular injection in P1 mice (Fig. 1A). As a control, we injected mice with either AAV9-GFP or PBS. After injections, mice were monitored for survival and behavior for up to 6 months.

**Figure 1.**
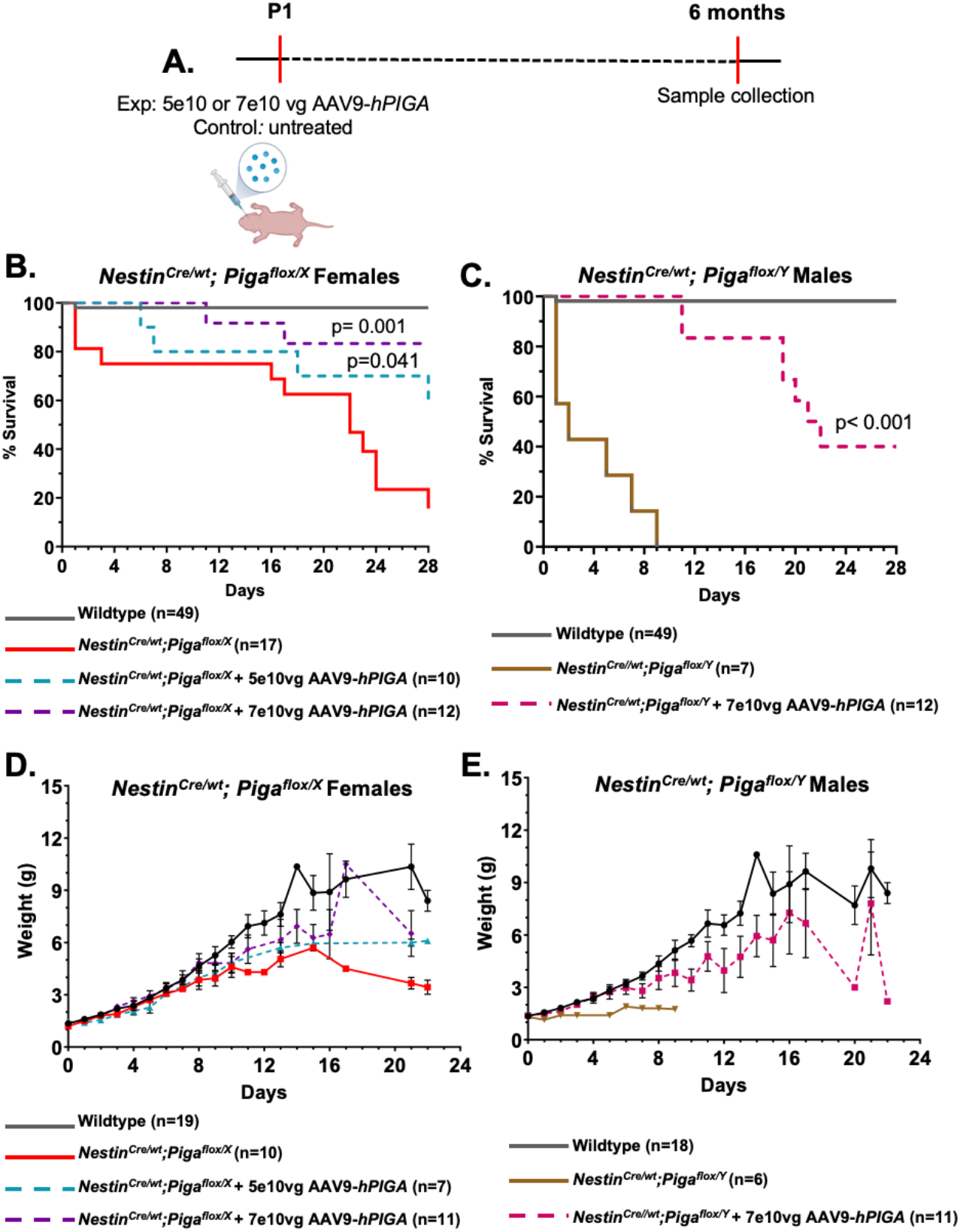
AAV9-*hPIGA* rescues survival in *NestinCre;Piga* females and males. **A)** AAV9-*hPIGA* injection and experimental design. **B)** Kaplan-Meier survival curves of wildtype, *Nestin*^*Cre/wt*^; *Piga*^*flox/X*^ females, *Nestin*^*Cre/wt*^; *Piga*^*flox/X*^ females treated with 5e10 viral genomes (vg) of AAV9-*hPIGA* (Mantel-Cox p=0.041) and *Nestin*^*Cre/wt*^; *Piga*^*flox/X*^ females treated with 7e10vg of AAV9-*hPIGA* (Mantel-Cox p=0.001). **C)** Kaplan-Meier survival curves of wildtype, *Nestin*^*Cre/wt*^; *Piga*^*flox/Y*^ males, and *Nestin*^*Cre/wt*^; *Piga*^*flox/Y*^ males treated with 7e10vg of AAV9-*hPIGA* (Mantel-Cox p<0.001). **D)** Body weights of wildtype, *Nestin*^*Cre/wt*^; *Piga*^*flox/X*^ females, *Nestin*^*Cre/wt*^; *Piga*^*flox/X*^ females treated with 5e10vg of AAV9-*hPIGA*, and *Nestin*^*Cre/wt*^; *Piga*^*flox/X*^ females treated with 7e10vg of AAV9-*hPIGA*. **E)** Body weights of wildtype, *Nestin*^*Cre/wt*^; *Piga*^*flox/Y*^ males, and *Nestin*^*Cre/wt*^; *Piga*^*flox/Y*^ males treated with 7e10vg of AAV9-*hPIGA*.

Wildtype mice showed a 98% survival rate at P28 (n=50, Mantel-Cox p<0.001). *Nestin*^*Cre/wt*^; *Piga*^*flox/X*^ females treated with a single injection of 5e10vg AAV9-*hPIGA* survive to P28 at significantly higher rates (n=10, Mantel-Cox p=0.041) than untreated *Nestin*^*Cre/wt*^; *Piga*^*flox/X*^ females (n=17). The survival rate of mice treated with 7e10vg of AAV9-*hPIGA* also improved at P28 compared to untreated *Nestin*^*Cre/wt*^; *Piga*^*flox/X*^ females (n=12, Mantel-Cox Test p=0.001, Fig. 1B). Fifty percent of *Nestin*^*Cre/wt*^; *Piga*^*flox/Y*^ males expire at P1 which made it difficult to robustly test this regimen in hemizygous male mice. The *Nestin*^*Cre/wt*^; *Piga*^*flox/Y*^ males that did survive to P1 were given a single dose of 7e10vg AAV-*hPIGA* and survived at significantly higher rates (n=12) than untreated *Nestin*^*Cre/wt*^; *Piga*^*flox/Y*^ males (n=6, Mantel-Cox p<0.001) which do not survive past P9 as previously reported (42) (Fig. 1C). We also noted an increase in body weight upon treatment with AAV9-*hPIGA* injection as surviving mice are significantly heavier than untreated mice (Table 1-2, Fig. 1D-E). These results reveal that a single injection of AAV9- *hPIGA* as a gene replacement therapy significantly increases survival and body weight in Piga mutant mice in a dose-dependent manner.

**Table 1.** Statistical analysis of *Nestin*^*Cre/wt*^; *Piga*^*flox/X*^ body weight data. ANOVA and Tukey’s multiple comparison analysis p-values for body weights in wildtype, AAV9-*hPIGA* treated and untreated *Nestin*^*Cre/wt*^; *Piga*^*flox/X*^ females per day. Grey cells highlight p-values <0.05.

| Days | wt vs<br>*Nestin*<sup>Cre/wt</sup>; *Piga*<sup>flax/x</sup> | wt vs *Nestin*<sup>Cre/wt</sup>; *Piga*<sup>flax/x</sup><br>+<br>5e10vg AAV9-*hPIGA* | wt vs<br>*Nestin*<sup>Cre/wt</sup>; *Piga*<sup>flax/x</sup> +<br>7e10vg AAV9-*hPIGA* | *Nestin*<sup>Cre/wt</sup>; *Piga*<sup>flax/x</sup> vs<br>*Nestin*<sup>Cre/wt</sup>; *Piga*<sup>flax/x</sup> +<br>5e10vg AAV9-*hPIGA* | *Nestin*<sup>Cre/wt</sup>; *Piga*<sup>flax/x</sup> vs<br>*Nestin*<sup>Cre/wt</sup>; *Piga*<sup>flax/x</sup> +<br>7e10vg AAV9-*hPIGA* | *Nestin*<sup>Cre/wt</sup>; *Piga*<sup>flax/x</sup> +<br>5e10vg AAV9-*hPIGA* vs<br>*Nestin*<sup>Cre/wt</sup>; *Piga*<sup>flax/x</sup> +<br>7e10vg AAV9-*hPIGA* | ANOVA<br>p-value |
| --- | --- | --- | --- | --- | --- | --- | --- |
| P0 | 0.637 | 0.893 | 0.985 | 0.444 | 0.634 | 0.996 | 0.489 |
| P1 | 0.937 | 0.655 | 0.846 | 0.972 | 0.999 | 0.888 | 0.614 |
| P2 | 0.998 | 0.653 | 0.862 | 0.940 | >0.999 | 0.887 | 0.649 |
| P3 | 0.750 | 0.277 | 0.945 | 0.999 | 0.617 | 0.254 | 0.19 |
| P4 | 0.999 | 0.822 | 0.871 | 0.971 | 0.922 | 0.446 | 0.515 |
| P5 | 0.987 | 0.488 | 0.986 | 0.806 | 0.943 | 0.407 | 0.449 |
| P6 | 0.940 | 0.940 | 0.999 | >0.999 | 0.972 | 0.972 | 0.903 |
| P7 | 0.837 | 0.924 | >0.999 | 0.998 | 0.886 | 0.950 | 0.802 |
| P8 | 0.851 | 0.912 | 0.990 | >0.999 | 0.770 | 0.844 | 0.713 |
| P9 | 0.608 | 0.772 | 0.907 | 0.995 | 0.829 | 0.944 | 0.583 |
| P10 | 0.466 | 0.617 | 0.254 | 0.997 | 0.996 | >0.999 | 0.218 |
| P11 | 0.364 | — | 0.457 | — | 0.757 | — | 0.295 |
| P12 | 0.306 | — | 0.175 | — | 0.873 | — | 0.145 |
| P13 | 0.341 | 0.570 | 0.595 | 0.982 | 0.889 | 0.991 | 0.294 |
| P15 | 0.525 | 0.386 | 0.216 | >0.999 | 0.993 | 0.997 | 0.21 |
| P17 | 0.207 | — | 0.609 | — | 0.623 | — | 0.226 |
| P21 | 0.373 | 0.365 | 0.931 | 0.981 | 0.338 | 0.351 | **0.024** |
| P22 | **0.003** | 0.054 | — | **0.025** | — | — | **0.003** |
| Overall | **0.011** | **0.034** | 0.239 | 0.994 | 0.554 | 0.754 | **0.009** |

To test if AAV9-*hPIGA* treatment leads to increased PIGA protein, we analyzed PIGA protein levels using immunohistochemistry. We observed that PIGA levels in Purkinje cells were increased, and cell bodies were again spatially organized in treated *Nestin*^*Cre/wt*^; *Piga*^*flox/X*^ females (n=4) and *Nestin*^*Cre/wt*^; *Piga*^*flox/Y*^ males (n=3) compared to untreated *Nestin*^*Cre/wt*^; *Piga*^*flox/X*^ (n=2) females at weaning (Fig. 2A-D). We previously reported that *Nestin* mediated *Piga* ablation causes reduced myelin in *Nestin*^*Cre/wt*^; *Piga*^*flox/X*^ mice (42). We observed recovery of myelin in brain sections of treated *Nestin*^*Cre/wt*^; *Piga*^*flox/X*^ females as early as weaning age (Fig.2E-H). Purkinje cell-specific PIGA levels in treated *Nestin*^*Cre/wt*^; *Piga*^*flox/X*^ females and *Nestin*^*Cre/wt*^; *Piga*^*flox/Y*^ males were sustained for at least 6 months and comparable to wildtype at that time point (n=3; Fig.2I-K). Additionally, western immunoblot of lysate from wildtype and treated *Nestin*^*Cre/wt*^; *Piga*^*flox/X*^ females tissue revealed no significant difference in PIGA levels (ANOVA p=0.935, Fig.2L, M, S1A). Myelin recovery was sustained for 6 months in treated *Nestin*^*Cre/wt*^; *Piga*^*flox/X*^ females and *Nestin*^*Cre/wt*^; *Piga*^*flox/Y*^ males (Fig.2N-P). We observed no qualitative differences in PIGA immunohistochemistry or behavioral outcomes in low or high dose of AAV9- *hPIGA* treated females. AAV9-*hPIGA* treatment results in long-term improvement in brain structure and development because of restored PIGA proteins levels and function.

**Figure 2.**
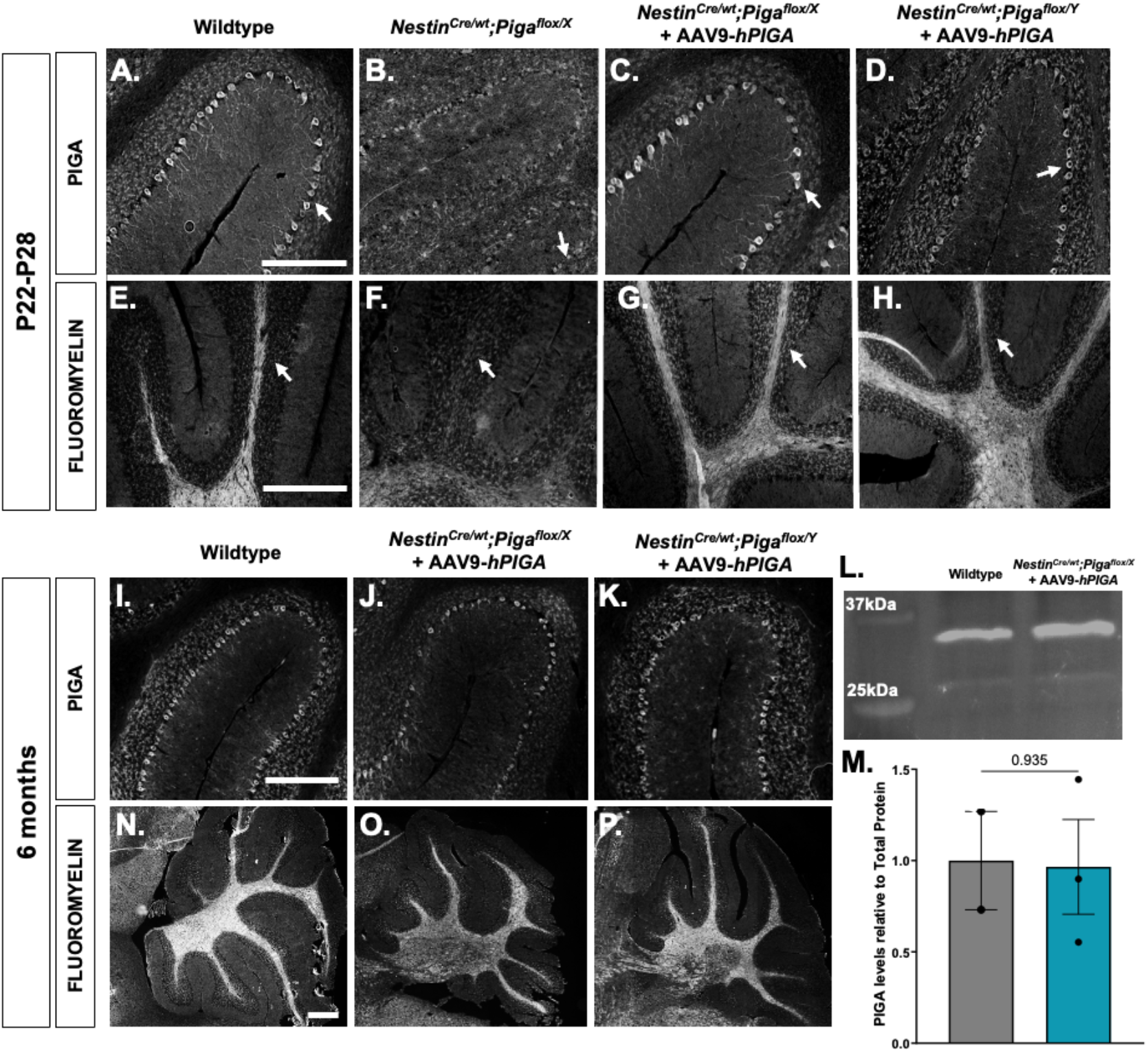
*NestinCre;Piga* animals injected with AAV9-*hPIGA* show restored PIGA and Myelin levels. PIGA levels and Purkinje cell organization (**A-D** scale bar = 250 µm) and fluoromyelin staining (**E-H** scale bar = 500 µm) in wildtype (A, E, n=4), *Nestin*^*Cre/wt*^; *Piga*^*flox/X*^ females (B, F, n=2), *Nestin*^*Cre/wt*^; *Piga*^*flox/X*^ AAV9-*hPIGA* treated females (C, G, n=4), and *Nestin*^*Cre/wt*^; *Piga*^*flox/Y*^ AAV9-*hPIGA* treated males (D, H, n=2) at P22-28 and at 6 months (**I-K, N-P**, scale bar = 500 µm). White arrows point to Purkinje cells positive for PIGA protein and putative fluoromyelin areas. (**L-M)** Protein blot and quantification of immunoblot PIGA levels at 6 months.

### Piga-associated ataxic phenotypes are reduced by AAV9-hPIGA gene replacement therapy

Having demonstrated an increase in survival and restoration of PIGA levels, we determined if PIGA gene replacement rescues the ataxia phenotype previously seen in *Nestin*^*Cre/wt*^; *Piga*^*flox/X*^ females. We scored hindlimb clasping on a scale from 0 (no clasping) to 4 (severe clasping lasting 10 seconds) of wildtype, untreated *Nestin*^*Cre/wt*^; *Piga*^*flox/X*^ females and *PIGA* treated *Nestin*^*Cre/wt*^; *Piga*^*flox/X*^ females at P21. Untreated *Nestin*^*Cre/wt*^; *Piga*^*flox/X*^ females exhibit severe hindlimb clasping and treated *Nestin*^*Cre/wt*^; *Piga*^*flox/X*^ females exhibit mild hindlimb clasping showing that AAV9-*hPIGA* treatment can partially rescue the hindlimb clasping phenotype (ANOVA p=0.004, Fig.3A-D). In the Rotarod assay, we observed that the average latency to fall in 5e10 vg AAV9-*hPIGA* treated *Nestin*^*Cre/wt*^; *Piga*^*flox/X*^ females (n=3) was comparable to that seen in wildtype females (n=4, 2 months, p=0.103; 4 months, p=0.887; and 6 months, p=0.764). The latency to fall in 7e10 vg AAV9-*hPIGA* treated *Nestin*^*Cre/wt*^; *Piga*^*flox/X*^ females (n=6) was also comparable to wildtype females (n=4, 2 months, p=0.679; 4 months, p=0.235; and 6 months, p=0.437, Fig.3E). We observed similar results in treated *Nestin*^*Cre/wt*^; *Piga*^*flox/Y*^ (n=2) and wildtype males (n=5, 2 months, p=0.940; 4 months, p=0.961; and 6 months, p>0.999, Fig.3F).

Finally, we measured body weight in wildtype and treated mutant mice and found that 5e10 vg AAV9-*hPIGA* treated *Nestin*^*Cre/wt*^; *Piga*^*flox/X*^ female body weights were slightly reduced (2 months, p=0.112; 4 months, p=0.570; and 6 months, p=0.113). Similarly, 7e10 vg AAV9- *hPIGA* treated *Nestin*^*Cre/wt*^; *Piga*^*flox/X*^ female body weights were slightly reduced (2 months, p=0.771; 4 months, p=0.472; and 6 months, p=0.062, Fig. 3G). Treated *Nestin*^*Cre/wt*^; *Piga*^*flox/Y*^ males body weights were comparable to wildtype males up to 6 months (2 months, p=0.541, 4 months, p=0.924; and 6 months, p=0.983, SFig.3H). We do note that the *Nestin*^*Cre/wt*^; *Piga*^*flox/Y*^ male studies are underpowered due to the significantly decreased survival in these animals from the injection time point onward. Overall, we demonstrate promising results that a singular injection of AAV9-*hPIGA* rescues the survival, neurological, and structural brain phenotypes in mice with CNS-specific ablations. These studies serve as a preclinical trial model for treatment in *PIGA* patients and could be a breakthrough treatment for patients with *PIGA* variant phenotypes, especially infant mortality.

**Figure 3.**
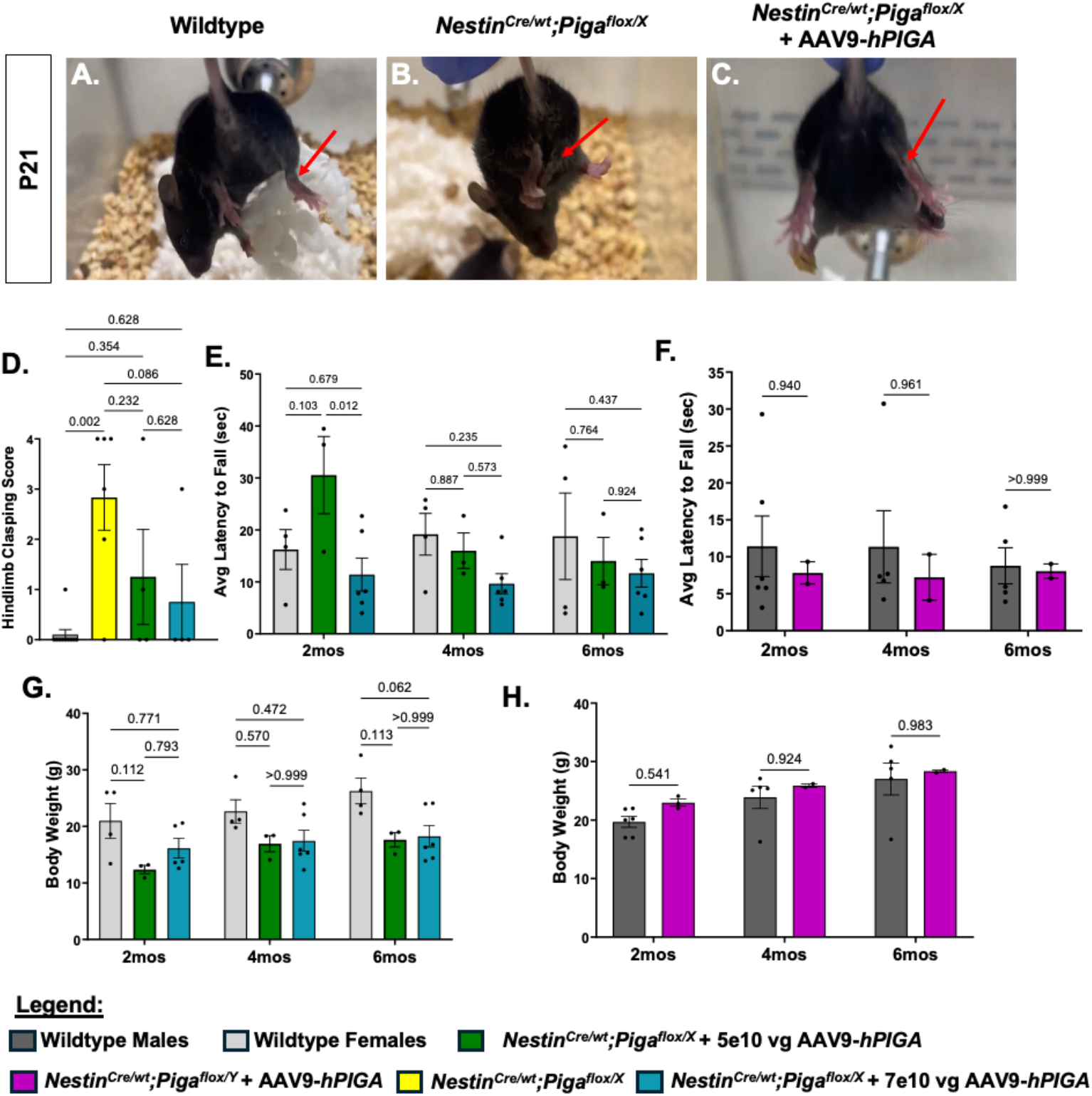
Reduced ataxia in treated *NestinCre;Piga* animals. **A)** Images of the degrees of hindlimb clasping in wildtype, **B)** *Nestin*^*Cre/wt*^; *Piga*^*flox/X*^ females and **C)** *Nestin*^*Cre/wt*^; *Piga*^*flox/X*^ females treated with AAV9-*hPIGA* at P21. **D)** Hindlimb clasping scoring (0=no clasping, 4= hindlimb clasped together) of wildtype, *Nestin*^*Cre/wt*^; *Piga*^*flox/X*^ females and *Nestin*^*Cre/wt*^; *Piga*^*flox/X*^ AAV9-*hPIGA* treated females at P21. **E-F)** Recorded Rotarod Performance Assay latency to fall in seconds and **G-H)** body weights for wildtype females, *Nestin*^*Cre/wt*^; *Piga*^*flox/X*^ females treated with AAV9-*hPIGA*, wildtype males, and *Nestin*^*Cre/wt*^; *Piga*^*flox/Y*^ males treated with AAV9-*hPIGA* at 2-, 4-, and 6-months post ICV injections.

### PIGA function directly affects the levels of GPI-anchored proteins

Finally, we wanted to identify which GPI-anchored proteins are affected by changes in *Piga* expression in the brain. We employed the alpha-toxin capture method to directly pull-down GPI-anchored proteins in brain lysates (61-65). We then observed the levels of GPI-anchored proteins to test the hypothesis that PIGA gene replacement therapy will rescue GPI-anchor proteins levels in mutant animal brains. We collected brain tissue from wildtype, untreated and treated *Nestin*^*Cre/wt*^; *Piga*^*flox/X*^ females, and untreated and treated *Nestin*^*Cre/wt*^; *Piga*^*flox/Y*^ males at P5 (4 animals per genotype per treatment). We used alpha-toxin to capture GPI-anchored proteins in a pull-down assay and liquid chromatography label free mass spectrometry to identify the abundance of captured proteins in each sample relative to total spectra. We compiled a list of putative GPI-anchored proteins that are expressed in the brain and are associated with neural phenotypes. The mass spectrometry results revealed that 31 of these 57 putative neuronal GPI-anchor proteins were present in our samples (listed in bold, SFig.2A). We noticed another 2,606 proteins identified between the samples showing that we precipitated complexes bound to GPI proteins, not just GPI-linked proteins (Sup.Table1).

We first wanted to investigate differentially downregulated alpha-toxin captured proteins in *Nestin*^*Cre/wt*^; *Piga*^*flox/X*^ female and *Nestin*^*Cre/wt*^; *Piga*^*flox/Y*^ male mutants (LOG2 fold change of <0 and p value < 0.1). *Nestin*^*Cre/wt*^; *Piga*^*flox/X*^ females had 30 unique differentially downregulated proteins whereas *Nestin*^*Cre/wt*^; *Piga*^*flox/Y*^ males had 83 unique differentially downregulated proteins (Fig.4A). These results revealed that more proteins are differentially downregulated in *Nestin*^*Cre/wt*^; *Piga*^*flox/Y*^ males, consistent with the more extreme phenotypes seen in males. Twelve proteins are commonly differently downregulated in both *Nestin*^*Cre/wt*^; *Piga*^*flox/X*^ females and *Nestin*^*Cre/wt*^; *Piga*^*flox/Y*^ males. These include the GPI-anchored proteins, LY6H, PRNP, RGMA, CDH13, and other proteins interacting with GPI-anchored proteins, POGZ, CORO1B, ERLIN2, EIF2B4, EPS8, SASH1, SMAD4, and RPS20 (Fig 4B).

**Figure 4.**
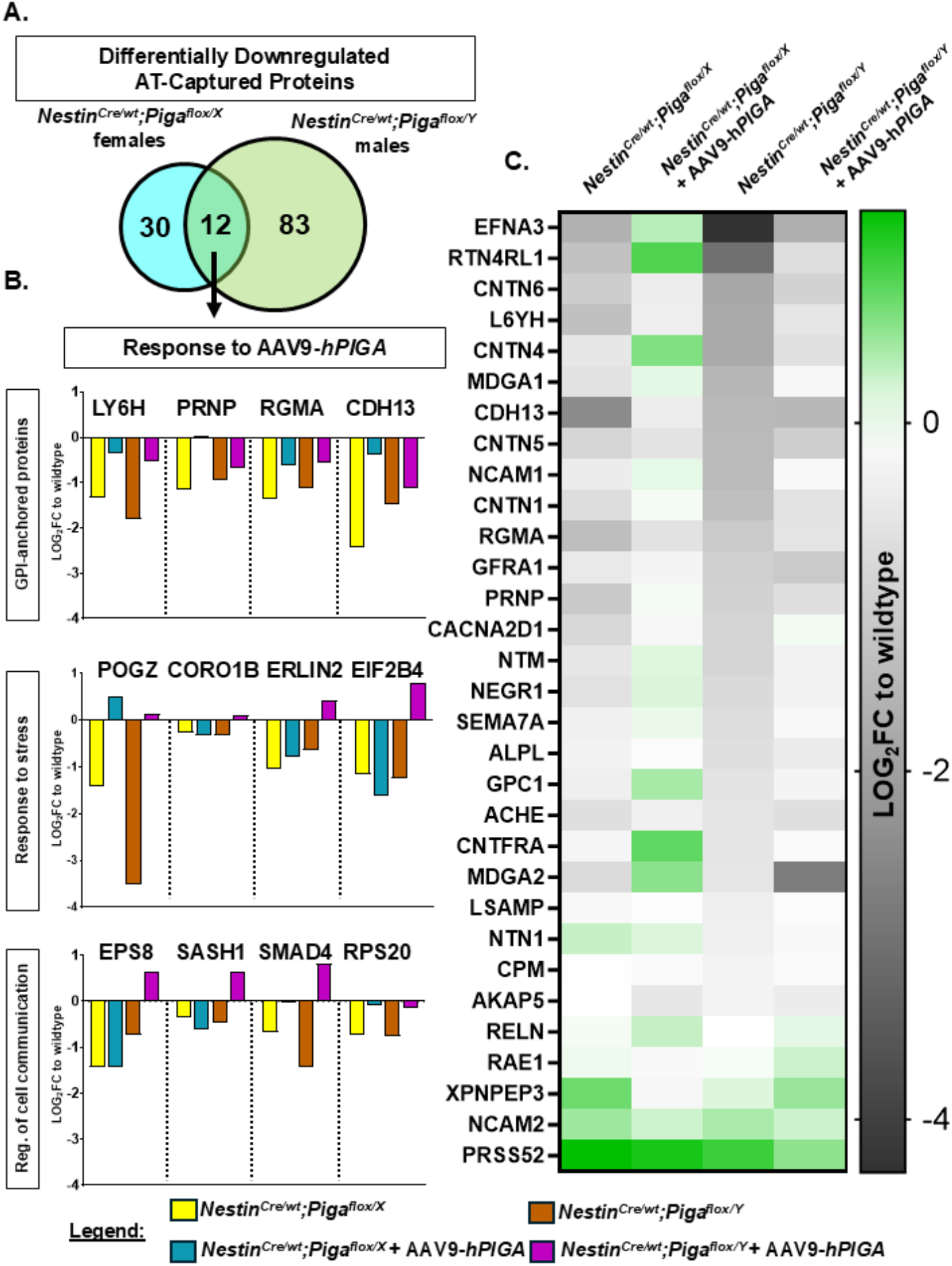
AAV9-*hPIGA* recovers GPI-anchored protein and GPI-anchor protein associated proteins in NestinCre;*Piga* mutants. **A)** Venn-Diagram of differentially downregulated proteins in *Nestin*^*Cre/wt*^; *Piga*^*flox/X*^ females and *Nestin*^*Cre/wt*^; *Piga*^*flox/Y*^ males. 12 proteins are common between male and female mutants. **B)** Protein levels response to AAV9-*hPIGA* depicted by the LOG2 fold change to wildtype levels in *Nestin*^*Cre/wt*^; *Piga*^*flox/X*^ females, *Nestin*^*Cre/wt*^; *Piga*^*flox/X*^ AAV9-*hPIGA* treated females, *Nestin*^*Cre*^*;Piga*^*flox/Y*^ males, and *Nestin*^*Cre/wt*^; *Piga*^*flox/Y*^ AAV9-*hPIGA* treated males of GPI-anchored, response to stress and Reg. of cell communication twelve common downregulated proteins. **C)** A heatmap showing the LOG2 fold change to wildtype levels of all GPI-anchored proteins in *Nestin*^*Cre/wt*^; *Piga*^*flox/X*^ females, *Nestin*^*Cre/wt*^; *Piga*^*flox/X*^ females treated with AAV9-*hPIGA, Nestin*^*Cre/wt*^; *Piga*^*flox/Y*^ males, and *Nestin*^*Cre/wt*^; *Piga*^*flox/Y*^ males treated with AAV9-*hPIGA*. n=4 animals per genotype at P5.

We then wanted to understand the effect of AAV9-*hPIGA* treatment on the twelve proteins downregulated in both types of mutants. A gene ontology analysis (66) suggested that POGZ, CORO1B, ERLIN2, and EIF2B4 were mostly associated with a response to stress biological process (67-76) and EPS8, SASH1, SMAD4, and RPS20 were mostly associated with the regulation of cell communication (77-88) (Table 3, SFig. 2B). GPI-anchored proteins LY6H, PRNP, RGMA, CDH13 increased with AAV9- *hPIGA* injections in both *Nestin*^*Cre/wt*^; *Piga*^*flox/X*^ females and *Nestin*^*Cre/wt*^; *Piga*^*flox/Y*^ males. Proteins related to response to stress, POGZ, CORO1B, ERLIN2, and EIF2B4 increased in AAV9- *hPIGA* treated *Nestin*^*Cre/wt*^; *Piga*^*flox/X*^ females *and Nestin*^*Cre/wt*^; *Piga*^*flox/Y*^ males with the largest response seen in POGZ levels. Proteins related to regulation of cell communication, EPS8, SASH1, SMAD4, and RPS20 all increased in AAV9-*hPIGA* treated *Nestin*^*Cre/wt*^; *Piga*^*flox/Y*^ males. Only SMAD4 and RPS20 increased with AAV9-*hPIGA* treated *Nestin*^*Cre/wt*^; *Piga*^*flox/X*^ females (Fig. 4B). We then specifically analyzed the 31 putative GPI-anchored proteins in the proteomic data to understand the response to gene therapy for all GPI-anchors. We observed reduced levels in the majority proteins in untreated *Nestin*^*Cre/wt*^; *Piga*^*flox/X*^ females and *Nestin*^*Cre/wt*^; *Piga*^*flox/Y*^ males (black and grey cells) than in AAV9-*hPIGA* treated *Nestin*^*Cre/wt*^; *Piga*^*flox/X*^ females and *Nestin*^*Cre/wt*^; *Piga*^*flox/Y*^ males (white and green cells, Fig.4C). As expected, we see a decrease in almost all of these proteins in mutant animals and almost universal increase in the treated animals. These results show that gene replacement therapy generally increases GPI-anchored proteins and their interacting molecules in *Nestin*^*Cre/wt*^; *Piga*^*flox/X*^ female and *Nestin*^*Cre/wt*^; *Piga*^*flox/Y*^ male mutants.

**Table 2.**
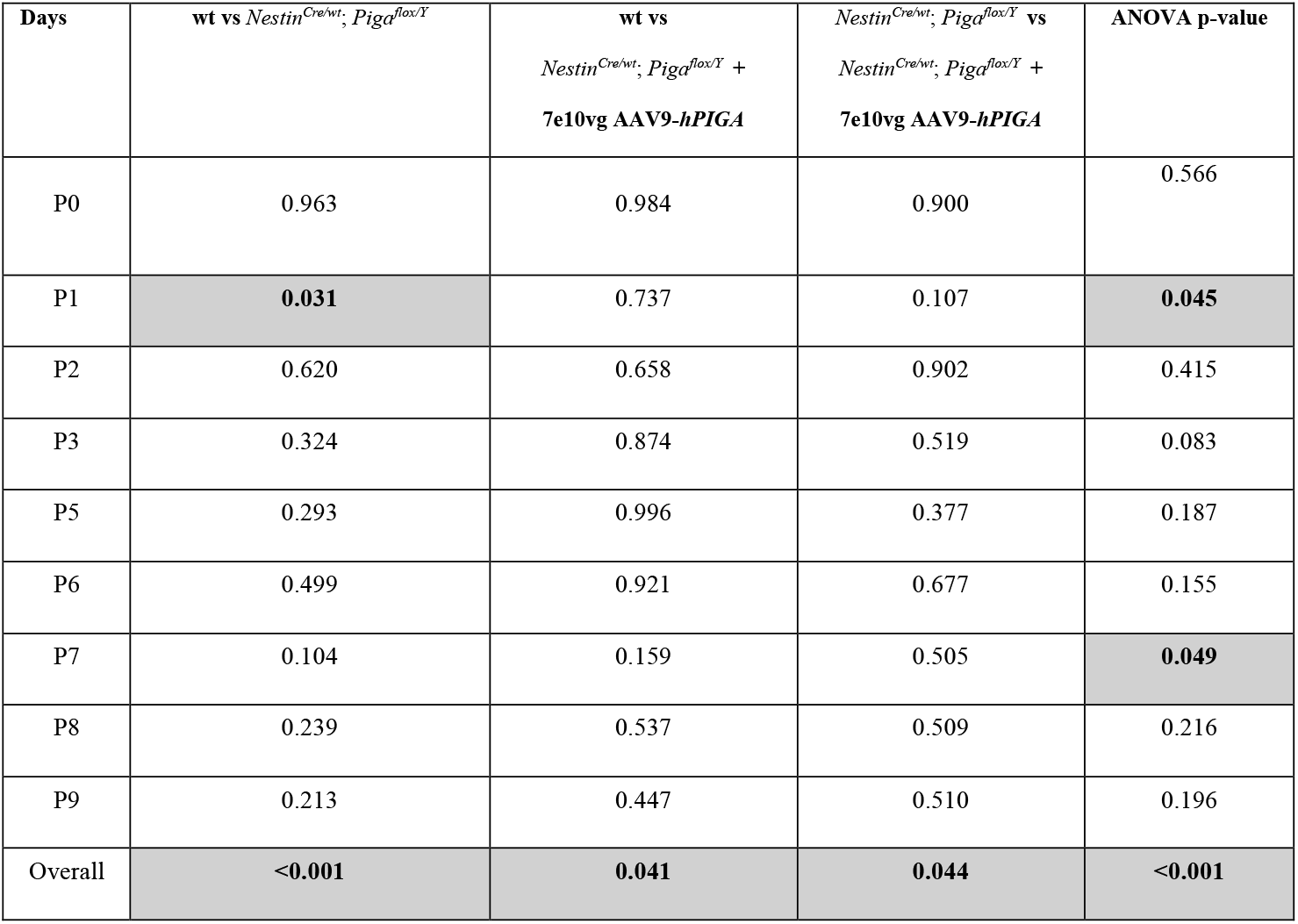
Statistical analysis of *Nestin*^*Cre/wt*^; *Piga*^*flox/Y*^ body weight data. ANOVA and Tukey’s multiple comparison analysis p-values for body weights in wildtype, AAV9-*hPIGA* treated and untreated *Nestin*^*Cre/wt*^; *Piga*^*flox/Y*^ males per day. Grey cells highlight p-values <0.05.

| Days | wt vs *Nestin*<sup>Cre/wt</sup>; *Piga*<sup>flax/Y</sup> | wt vs<br>*Nestin*<sup>Cre/wt</sup>; *Piga*<sup>flax/Y</sup> +<br>7e10vg AAV9-*hPIGA* | *Nestin*<sup>Cre/wt</sup>; *Piga*<sup>flax/Y</sup> vs<br>*Nestin*<sup>Cre/wt</sup>; *Piga*<sup>flax/Y</sup> +<br>7e10vg AAV9-*hPIGA* | ANOVA p-value |
| --- | --- | --- | --- | --- |
| P0 | 0.963 | 0.984 | 0.900 | 0.566 |
| P1 | **0.031** | 0.737 | 0.107 | **0.045** |
| P2 | 0.620 | 0.658 | 0.902 | 0.415 |
| P3 | 0.324 | 0.874 | 0.519 | 0.083 |
| P5 | 0.293 | 0.996 | 0.377 | 0.187 |
| P6 | 0.499 | 0.921 | 0.677 | 0.155 |
| P7 | 0.104 | 0.159 | 0.505 | **0.049** |
| P8 | 0.239 | 0.537 | 0.509 | 0.216 |
| P9 | 0.213 | 0.447 | 0.510 | 0.196 |
| Overall | **<0.001** | **0.041** | **0.044** | **<0.001** |

**Table 3.**
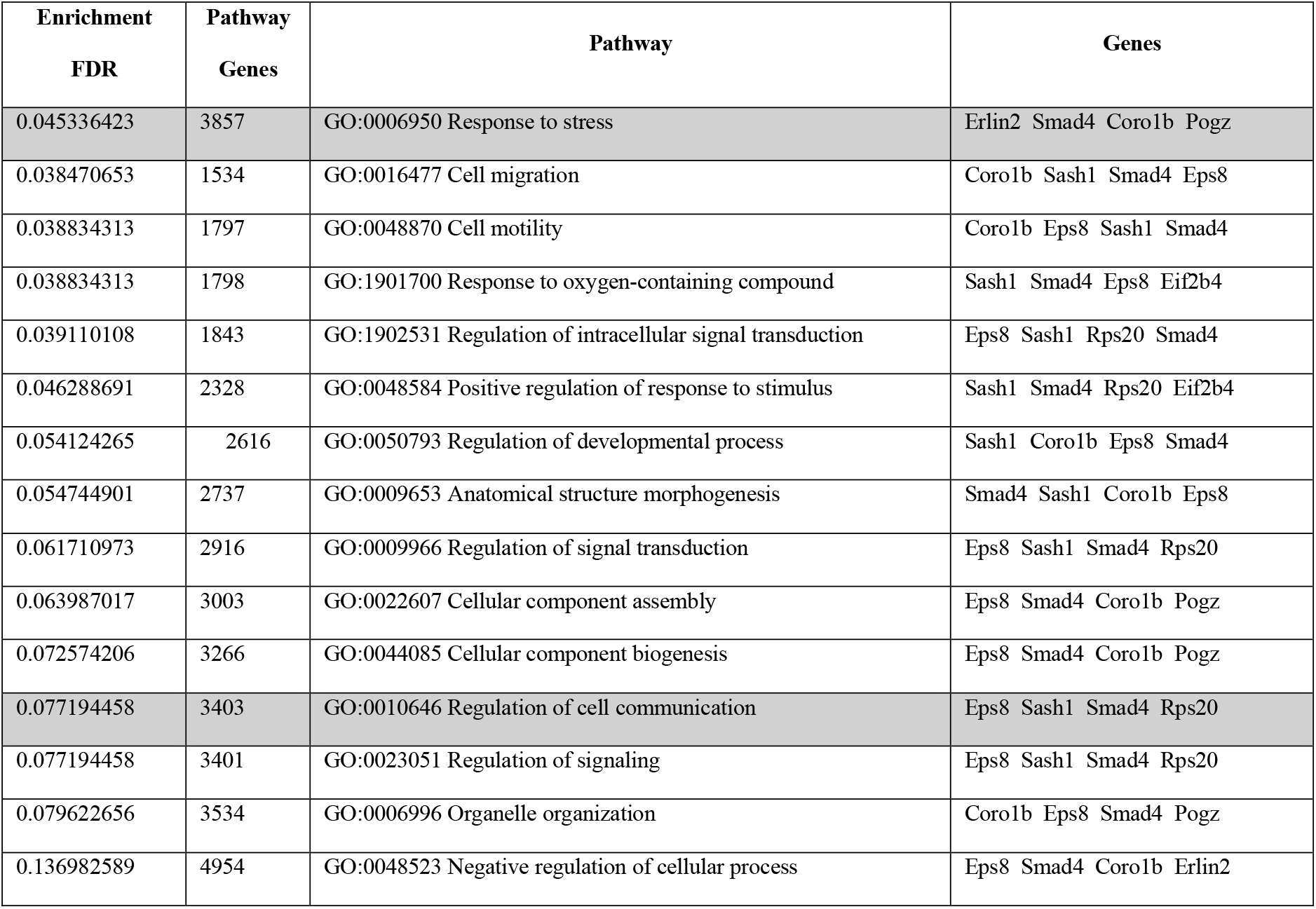
Gene Ontology of GPI-anchored protein partners. Gene Ontology (GO) analysis with Enrichment FDR, the number genes in each GO Pathway, GO term identification, and the downregulated GPI-anchored binding partners associated with each GO term from ShinyGO 0.85 database. Grey cells are GO term in response to stress genes and regulation of cell communication.

| Enrichment<br>FDR | Pathway<br>Genes | Pathway | Genes |
| --- | --- | --- | --- |
| 0.045336423 | 3857 | GO:0006950 Response to stress | Erlin2 Smad4 Coro1b Pogz |
| 0.038470653 | 1534 | GO:0016477 Cell migration | Coro1b Sash1 Smad4 Eps8 |
| 0.038834313 | 1797 | GO:0048870 Cell motility | Coro1b Eps8 Sash1 Smad4 |
| 0.038834313 | 1798 | GO:1901700 Response to oxygen-containing compound | Sash1 Smad4 Eps8 Eif2b4 |
| 0.039110108 | 1843 | GO:1902531 Regulation of intracellular signal transduction | Eps8 Sash1 Rps20 Smad4 |
| 0.046288691 | 2328 | GO:0048584 Positive regulation of response to stimulus | Sash1 Smad4 Rps20 Eif2b4 |
| 0.054124265 | 2616 | GO:0050793 Regulation of developmental process | Sash1 Coro1b Eps8 Smad4 |
| 0.054744901 | 2737 | GO:0009653 Anatomical structure morphogenesis | Smad4 Sash1 Coro1b Eps8 |
| 0.061710973 | 2916 | GO:0009966 Regulation of signal transduction | Eps8 Sash1 Smad4 Rps20 |
| 0.063987017 | 3003 | GO:0022607 Cellular component assembly | Eps8 Smad4 Coro1b Pogz |
| 0.072574206 | 3266 | GO:0044085 Cellular component biogenesis | Eps8 Smad4 Coro1b Pogz |
| 0.077194458 | 3403 | GO:0010646 Regulation of cell communication | Eps8 Sash1 Smad4 Rps20 |
| 0.077194458 | 3401 | GO:0023051 Regulation of signaling | Eps8 Sash1 Smad4 Rps20 |
| 0.079622656 | 3534 | GO:0006996 Organelle organization | Coro1b Eps8 Smad4 Pogz |
| 0.136982589 | 4954 | GO:0048523 Negative regulation of cellular process | Eps8 Smad4 Coro1b Erlin2 |

## Discussion

Congenital anomalies due to inherited GPI deficiencies can include numerous structural brain defects and neurological symptoms such as intellectual disability, developmental delay, hypomyelination, and cerebellar hypoplasia (10, 12, 25, 89). Some severe cases can even exhibit microcephaly and infant mortality (10, 17, 18, 21-24, 26-28, 32, 90). Our prior studies involving a central nervous system deletion of *Piga* (Nestin-Cre) in murine models replicated many human symptoms, thereby supporting the validity of this model for human *PIGA* variants (42). Advances in AAV9-mediated gene-replacement therapies underscore the potential to use these mouse *Piga* models to evaluate treatment efficacy. This study demonstrates that a single administration of AAV9-*hPIGA* can rescue multiple structural brain defects and neurological phenotypes in mutant mice and restores the levels of specific downstream neuronal GPI-anchored proteins and their associated partners.

Previous attempts to treat neuronal *Piga* phenotypes involved delivering the human PIGA transcript via an AAV-PHPeB virus and intravenous injections. These efforts resulted in several weeks of improved survival and reduced ataxia symptoms in injected mutants compared to untreated ones. Notably however, systemic injections led to liver tumors in animals within a year of injection (40). The are significant differences between this study and the one presented here. We used AAV9, which has high affinity for the mammalian brain, and administered it directly into the brain via intracerebroventricular injections, which likely explains the contrasting results. In our study, we observed substantial improvements in survival up to 6 months, along with higher PIGA levels and proper localization, and reduced ataxia symptoms in mutant treated animals compared to untreated mutants. Furthermore, we found that AAV9-*hPIGA* dosing is crucial for achieving better treatment outcomes. The proteomic data generated in this study is different from the previous attempt at gene therapy for *Piga*-related neurological deficiencies where there was no significant increase in GPI-anchored proteins seen in the treated males (40). However, our approach took advantage of the specificity of the alpha-toxin-mediated GPI protein capture to measure GPI-anchored protein levels in treated and untreated mutant mice. We hypothesize that the route of injections, viral packaging, and the alpha-toxin capture could contribute to the power of the gene replacement therapy and selection of proteins in this study.

There are about 150 known GPI-anchored proteins, and approximately 60 of those have an established role in neural development (13, 91). The proteomic data show that treatment in both *Nestin*^*Cre/wt*^; *Piga*^*flox/X*^ females and *Nestin*^*Cre/wt*^; *Piga*^*flox/Y*^ males recovered many GPI-anchored proteins such as the CONTACTIN proteins, LY6H, and NTM (Fig.4C, S2B-C). However, proteins such as MDGA2, AKAP5, ACHE, and GFRA1 were still found at low levels after treatment in *Nestin*^*Cre/wt*^; *Piga*^*flox/Y*^ male animals (S2C-D). These results suggest that these proteins could be essential for survival and an indicator of treatment outcomes in neurodevelopmental *PIGA* associated dysfunction (49, 92-98). We also identified putative non-GPI-anchored proteins, which we concluded to be GPI-anchored protein binding partners. We also question whether we identified molecules that have not yet been classified as GPI-anchored proteins. The findings concerning GPI-anchored proteins partners related to response to stress and cell communication found in our dataset were also interesting because they address downstream effects of unmodified GPI-anchor proteins in the brain (72). POGZ, a response to stress protein, had one of the largest increases in treated animals. Interestingly, variants in *POGZ* cause intellectual disability, microcephaly, facial dysmorphism, and epilepsy, much like *PIGA* variants (18, 21, 32, 99).

However, there is no evidence for direct interactions between POGZ and PIGA and further studies would be needed to understand the connection between PIGA and POGZ activity. Proteins in this dataset could foster further exploratory investigations and expand our knowledge of GPI-anchored proteins. Also, identifying GPI-anchored proteins and their partners affected by *Piga* expression can serve as potential targets to complement gene-replacement therapy.

*Nestin*^*Cre/wt*^; *Piga*^*flox/X*^ females had better treatment outcomes than *Nestin*^*Cre/wt*^; *Piga*^*flox/Y*^ males which is likely due to random X-inactivation of *Nestin*^*Cre/wt*^; *Piga*^*flox/X*^ female mice leading to some retained expression of wildtype *Piga*. Precisely which cells retain *Piga* expression likely has a marked effect on the phenotype. The results from this study suggests that *Nestin*^*Cre/wt*^; *Piga*^*flox/Y*^ males may require a more powerful *PIGA* expression to rescue the complete loss of *Piga* expression. Incredibly, about 40% of *Nestin*^*Cre/wt*^; *Piga*^*flox/Y*^ males survived to P28 in the current treatment strategy with AAV9-*hPIGA* which was significantly higher than untreated mutant males that died by P9. These few mice that did survive were used for further testing and analysis, but the neonatal lethality of this genotype left some experiments somewhat underpowered by the 6-month time point. Nevertheless, our gene therapy study sets a precedent to treat both mild and severe cases of *PIGA* variants in humans.

While the current gene therapy approach effectively rescues many phenotypes in the developing brain, there is still work to be done to adapt this treatment for human *PIGA* variants. We are exploring ways to improve survival outcomes by administering earlier injections and/or increasing the dose of AAV9-*hPIGA*, which could enhance the power of long-term studies. Toxicology studies in both wild-type and mutant animals are essential to ensure the treatment’s safety. Additionally, most patients identified to date have a missense variant leading to a hypomorphic allele, unlike the knockout alleles used in these studies. Treating hypomorphic allele models will be the next step to fully assess the effectiveness of this gene replacement therapy in animal trials and its potential for human application. Here, we developed a therapeutic framework that will hopefully be a breakthrough for inherited GPI deficiency patients.

## Methods

### Animal husbandry

Mice were housed in a vivarium with a 12-hour light cycle with food and water *ad libitum*, and mouse lines were maintained on a C57Bl6/J background. *Piga*^*flox*^ (*B6*.*129-Piga*^*tm1*^, #RBRC06211) (100), previously generated by Taroh Kinoshita and Junji Takeda, were obtained from the RIKEN repository and genotyped with the following primers: Forward-ACCTCCAAAGACTGAGCTGTTG, Reverse-CCTGCCTTAGTCTTCCCAGTAC, and Forward LoxP site-TGTGGGTTTCAGTTCATTTCAGA. *B6*.*Cg-Tg(Nes-cre)*^*1Kln/J*^ *or Nestin-Cre* mice (Jackson labs #003771) (101) were genotyped for the *Cre* transgene using the following primers: Forward-GCGGTCTGGCAGTAAAAACTATC and Reverse-GTGAAACAGCATTGCTGTCACTT. Progeny of the *Piga*^*flox*^ and *Nestin*-Cre crosses were euthanized at the proper humane endpoint or until the end of the experiment.

#### Sex as a biological variable

*Piga* is X-linked in mouse and human. Therefore, in this study, we address AAV9-*hPIGA* treatment in male and female mice, as the *Piga* mutation exhibits sex specific-phenotypes. Sex of the mice was determined by the *Kdm5* gene the following primers: Forward-TGAAGCTTTTGGCTTTGAG and Reverse-CCGCTGCCAAATTCTTTGG.

### Virus Production

A human *PIGA* cDNA clone was obtained from Origene (Rockville, MD, USA) and subcloned into an AAV production vector under the control of a Chicken beta actin CMV promoter (102). Self-complementary AAV9.CMVCBA.*hPIGA* was produced by SAB Tech (Philadelphia, PA, USA) and Andelyn (Columbus, OH, USA) according to their established protocols. Purity and titer of the virus was assessed by silver staining and digital droplet (dd)PCR analysis, respectively.

### Intracerebroventricular injection of scAAV9.CBA.hPIGA

At P1, pups were genotyped and randomly assigned to groups of either GFP/PBS as a control or scAAV9.CBA.*hPIGA* treatment. The *Nestin*^*Cre/wt*^; *Piga*^*flox/X*^ and *Nestin*^*Cre/wt*^; *Piga*^*flox/Y*^ mice received a single injection at P1 with 5 µL intracerebroventricular (ICV) injections from a Hamilton syringe of either PBS or scAAV9.CBA.*hPIGA* at a dose of 5e10^10^ vg/animal or 7e10^10^ vg/animal, following hypothermia sedation. In addition to the treatment of *Nestin*^*Cre/wt*^; *Piga*^*flox/X*^ and *Nestin*^*Cre/wt*^; *Piga*^*flox/Y*^ animals, wildtype littermates were used as controls for treatment groups. All mice were monitored on a heating pad until fully recovered and returned to the dam.

### Immunostaining

Brains were collected and fixed in 4% paraformaldehyde for 24–48 hours. After washing in 30% sucrose, they were embedded in O.C.T. compound overnight or until they floated. The brains were then sectioned at 10 µm thickness using a cryostat. Frozen sections were heated at 42°C on a heat block for 10-15 minutes. Antigen retrieval was performed with citrate buffer followed by blocking with 4% normal goat serum. Slides were incubated in primary rabbit anti-PIGA (Proteintech #13679-1-AP) antibody at a 1:200 dilution overnight at 4°C. Afterwards, sections were incubated with 1:500 dilution of secondary Alexafluor 555-congugated goat anti-rabbit (Invitrogen # A21428) antibody or 1:300 dilution of conjugated antibody FluoroMyelin Green (Invitrogen #F34651) at room temperature. Sections were counterstained with DAPI and imaged with the Zeiss Apotome Fluorescence Microscope.

### Rotarod Motor Coordination Performance

Rotarod Motor Coordination testing was performed in the NCH Animal Behavior Core. Researchers were blinded to the animals’ genotypes. The Rotarod apparatus (San Diego Instruments) was used with the SDI software, following the 10 to 50 RPM (revolutions per minute) protocol for mice. The apparatus has four chambers, and up to four mice can be tested simultaneously. The photobeam sensor detects and records the time and distance each mouse travels before falling from the rod. The test was allowed to run for 300 s. The test starts at 10 RPM for 10 s and increases to 20 RPM for 10 s, then 30 RPM for 10 s, and finally 40 RPM for 10 s and 50 for 10 s. The latency to fall in seconds were recorded automatically and the mice were allowed to rest for 5 mins before each successive trial. A test trial followed by 3 recorded trials were performed sequentially on each mouse.

### Western immunoblotting

Brains were dissected and lysed in 1 mL RIPA buffer with Protease inhibitor. Lysate protein concentration was determined by BCA assay, and electrophoresis was performed on a 4-15% Tris-glycine stain-free precast gels (BioRad # 4568083) with All-Blue Protein Standard. Protein was transferred to a PVDF membrane via the Trans-Blot Turbo Transfer, blocked in BioRad Blocking buffer. Primary rabbit anti-PIGA (Proteintech #13679-1-AP) antibody were diluted to 1:1000 and incubated overnight at 4°C. Membranes were washed with PBST and incubated for 1 hour using the secondary antibody StarBright Blue 700 goat anti-rabbit (BioRad #12005866) at a 1:25000 dilution and visualized on a BioRad ChemiDoc imaging system. Relative protein concentration was determined by normalizing the protein of interest signal to total stain-free signal in Image Studio Lite Ver 5.2.

### Alpha Toxin Capture

The full protocol to produce and capture GPI-anchored proteins using alpha toxin capture was followed as described by Huang and Park for the proteomic analysis presented in this study (65).

### Mass Spectrometry

Beads were washed with 50 mM ammonium bicarbonate (100 µL each time) three times. Each time, the supernatant was kept and pooled. After the third wash, 5 µL of DTT (5 µg/µL in 50 mM ammonium bicarbonate) is added, and the sample is incubated at 56°C for 15 min. After the incubation, 5 µL of iodoacetamide (15 mg/mL in 50 mM ammonium bicarbonate) is added and the sample is kept in dark at room temperature for 30 min. Sequencing grade-modified trypsin (Promega, Madison WI) prepared in 50 mM ammonium bicarbonate and 1 µg of trypsin was added to the sample; the reaction was carried on at 37°C for overnight. The reaction is quenched the next morning by adding 10% Formic Acid for acidification. The supernatant was removed to concentrate proteins for LC/MSMS analysis.

Nano-liquid chromatography-nanospray tandem mass spectrometry (Nano-LC/MS/MS) of protein identification was performed on the Thermo Scientific orbitrap Eclipse mass spectrometer equipped with a nanospray FAIMS Pro™ Sources operated in positive ion mode. Samples (4 µL) were separated on an easy spray nano column (Pepmap™ RSLC, C18 3 µ 100A, 75 µm X250mm Thermo Scientific) using a 2D RSLC HPLC system from Thermo Scientific. Each sample was injected into the µ-Precolumn Cartridge (Thermo Scientific) and desalted with 0.1% Formic Acid in water for 5 minutes. The injector port was then switched to inject, and the peptides were eluted from the trap onto the column. Mobile phase A was 0.1% Formic Acid in water and acetonitrile (with 0.1% formic acid) was used as mobile phase B. Flow rate was set at 300 nL/min. mobile phase B was increased from 2% to 16% in 105 min and then increased from 16-25% in 10 min and again from 25-85% in 1 min and then kept at 95% for another 4 min before being brought back quickly to 2% in 1 min. The column was equilibrated at 2% of mobile phase B (or 98% A) for 15 min before the next sample injection.

MS/MS data was acquired with a spray voltage of 1.95 KV and a capillary temperature of 305°C is used. The scan sequence of the mass spectrometer was based on the preview mode data dependent TopSpeed™ method: the analysis was programmed for a full scan recorded between *m/z* 375-1500 and a MS/MS scan to generate product ion spectra to determine amino acid sequence in consecutive scans starting from the most abundant peaks in the spectrum in the next 3*1 seconds. To achieve high mass accuracy MS determination, the full scan was performed at FT mode and the resolution was set at 120,000 with internal mass calibration. Three compensation voltage (cv=- 40, –60 and –80v) were used for samples acquisition. The AGC Target ion number for FT full scan was set at 4 x 10^5^ ions, maximum ion injection time was set at auto and micro scan number was set at 1. MSn was performed using HCD in ion trap mode to ensure the highest signal intensity of MSn spectra. The HCD collision energy was set at 30%. The AGC Target ion number for ion trap MSn scan was set at 1.0E4 ions, maximum ion injection time was set at auto and micro scan number was set at 1. Dynamic exclusion is enabled with a repeat count of 1 within 20s and a low mass width and high mass width of 10ppm.

The resulting mfg.files generated from the samples were filtered using Mascot Daemon by Matrix Science version 2.7.0 (Boston, MA) against appropriate databases using the proteome discoverer (Thermo Fisher Scientific) platform. The mass accuracy of the precursor ions was set to 10 ppm, accidental pick of 1 13C peaks was also included into the search. The fragment mass tolerance was set to 0.5 Da. Considered variable modifications were oxidation (Met), deamidation (N and Q) and carbamidomethylation (Cys) is considered as fixed modification. Four missed cleavages for the enzyme were permitted. A decoy database was also searched to determine the false discovery rate (FDR) and peptides were filtered according to the FDR. The significance threshold was set at p<0.05 and bold red peptides is required for valid peptide identification. Proteins with less than 1% FDR as well as a minimum of 2 significant peptides detected were considered as valid proteins.

### Statistical analysis

Statistical analysis and graphs were generated using GraphPad Prism (GraphPad Software, San Diego, CA). Data presented in bar graphs and scatter plots are shown as the median with Standard Error of the Mean (SEM). Survival analysis was conducted with a log-rank Mantel-Cox test. Rotarod analysis was performed using a one-way ANOVA followed by Tukey’s multiple comparison test. Gene ontology analysis of the proteomics data was conducted with ShinyGO 0.85 (http://bioinformatics.sdstate.edu/go/). P-values displayed in bar graphs are the results of multiple comparisons testing unless otherwise specified.

## Acknowledgments

We want to thank the Stottmann laboratory members for their continuous counsel throughout this study. We would like to thank Dr. Park for providing the alpha-toxin plasmid and for the discussion regarding the proteomic portion of this study. We acknowledge resources from the Campus Chemical Instrumentation Center Mass Spectrometry and Proteomics Facility and the OSU Comprehensive Cancer Center (OSUCCC) Proteomics Shared Resource (PSR), The Ohio State University. This facility is supported in part by funding from OSU’s Enterprise for Research, Innovation and Knowledge and NCI grant P30 CA01605. Lastly, we would like to thank the funding sources from Nationwide Children’s Hospital Recruitment Funds, Abigail Wexner Research Institute Postdoctoral Diversity in Academia award, and the CDG Care Foundation, who supported the AAV9 gene therapy portion of this study.

## Author contributions

**Conceptualization**: R.W.S., K.M., T.K., Y.M., J.L.W; **Methodology:** J.L.W., R.W.S., K.M.; **Validation:** J.L.W., J.M.L. R.W.S.; **Formal analysis**: J.L.W.; **Investigation**: J.L.W.; **Resources**: R.W.S., K.M., S.L.; **Writing – original draft:** J.L.W; **Writing – review & editing:** J.L.W., S.L., T.K., K.M., R.W.S; **Visualization:** J.L.W; **Supervision**: R.W.S.; **Project administration:** R.W.S.; **Funding acquisition**: R.W.S., J.L.W., K.M. **Study Approval** The Nationwide Children’s Hospital IACUC committee (IACUC: AR21-00067) approved animal use and intracerebroventricular injection protocols for this study.

## Data Availability

## Lead contact

Further information and requests for resources and reagents should be directed to and will be fulfilled by the lead contact, Rolf Stottmann

## Materials availability

Materials will be fulfilled by the lead contact upon request

## Data and code availability

All data in this paper will be shared by the lead contact upon request.

No new code was produced in this paper Any additional information about the data acquired in this paper for further analysis is available from the lead contact upon request.

## Notes

**Conflict of Interest Statement.** The authors have declared that no conflict of interest exists.

### Competing Interest Statement

The authors have declared no competing interest.

