## Supplemental Data for "Gene replacement therapy for Piga GPI-anchor deficiency in the developing nervous system"

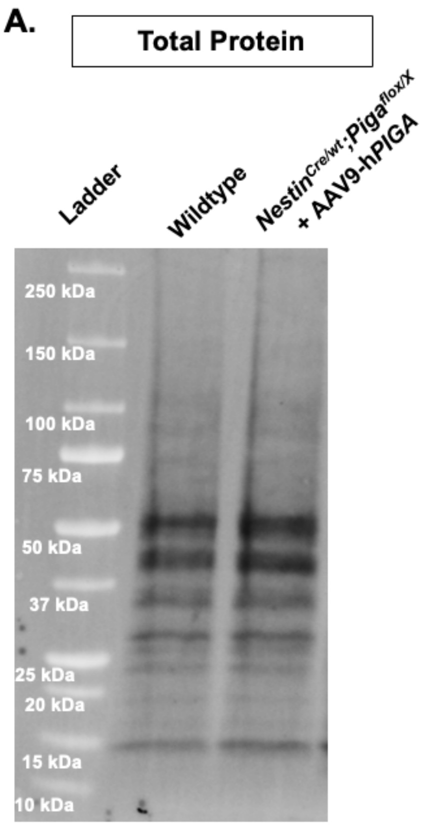


**SFigure 1- A)** Total protein stain-free blot for PIGA quantifications.

A.

| Putative Neuronal GPI- Anchored Proteins | |
|---|---|
| **AKAP5** | LYPD1 |
| **ALPL** | LYPD6 |
| CD59A | **MDGA1** |
| **CDH13** | **MDGA2** |
| CD59B | **NCAM1** |
| CNTFRA | **NCAM2** |
| **CNTN1** | **NEGR1** |
| CNTN2 | NRN1 |
| CNTN3 | **NTNG1/NTN1** |
| **CNTN4** | NTNG2 |
| **CNTN5** | **NTM** |
| **CNTN6** | OMGP |
| CNTNAP1 | OTOA |
| DAF1 | **PRION** |
| **EFNA3** | PSCA |
| ENPP6 | R4RL1 |
| GAS-1 | R4RL2 |
| GFRa1 | **RAE1A** |
| **GFRa2** | RAE1B |
| GFRa3 | RAE1C |
| **GPC1** | **RGMA** |
| GPC4 | RGMB |
| HYAL2 | **RTN4RL1** |
| IGS21 | **PRSS52** |
| ITL1A | **SEMA7A** |
| **LSAMP** | SPRN |
| LY6A | TDGF1 |
| **LY6H** | uPAR/Plaur |
| LY6I | XPP2 |
| LYNX1 | **XPNPEP3** |

Bolded GPI-anchored proteins found in mass spec data

C.

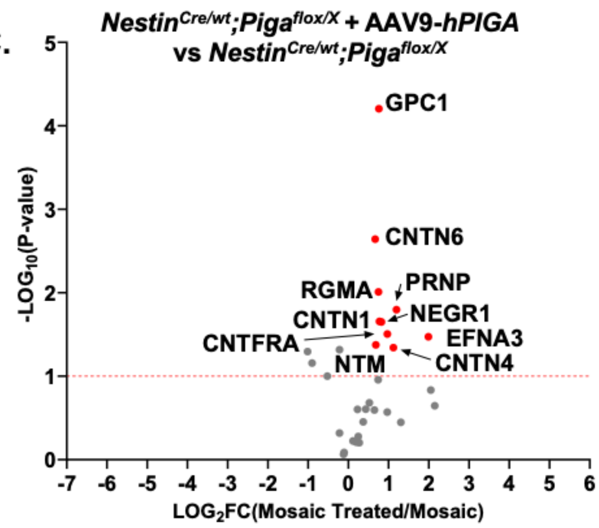


D.

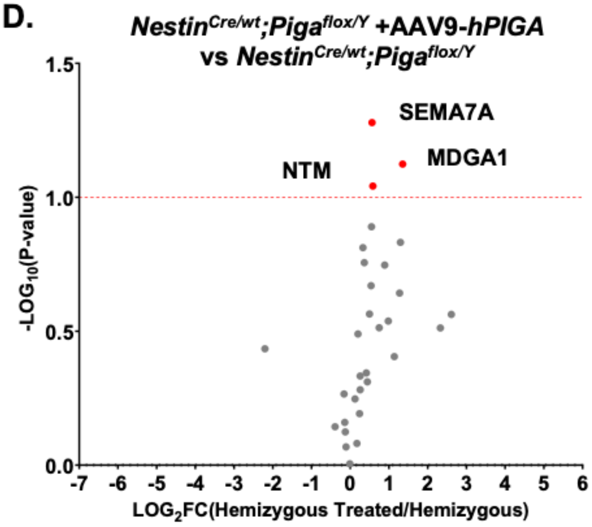


B.

Gene Ontology: biological processes of differentially downregulated proteins

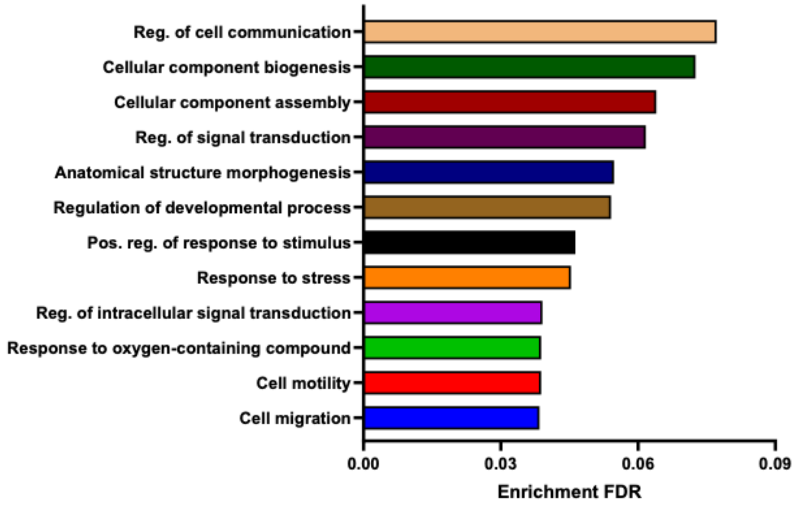

**SFigure 2 -** GPI-anchored protein levels in respons to AAV9-*hPIGA* between *NestinCre; Piga* females and males. **A)** Table of Putative Neural GPI-anchored proteins. Bolded proteins are found in experimental samples in this study. **B)** Gene Ontology analysis Enrichment False Discovery Rate (FDR) of downregulated non-GPI-anchored proteins. **C)** Volcano plot comparing the GPI-anchored proteins of $Nestin^{Cre/wt}$; $Piga^{flox/X}$ AAV9-*hPIGA* treated females and $Nestin^{Cre/wt}$;$Piga^{flox/X}$ untreated females and (**D**) comparing $Nestin^{Cre/wt}$;$Piga^{flox/Y}$ AAV9-*hPIGA* treated males and $Nestin^{Cre/wt}$;$Piga^{flox/Y}$ untreated males. Proteins in red are above the red dotted are differentially up- or downregulated. Proteins named are upregulated with *AAV9-hPIGA* injections (LOG2 fold change >0 and -LOG10pvalue >1).
